# Staphylococcal Internalization into Osteoblasts: A Partially Conserved Mechanism Across the Genus

**DOI:** 10.1101/2025.02.13.638029

**Authors:** Deborah M Crepin, Melanie Bonhomme, Allison Faure, Clara Sinel, Marine Bergot, Virginie Dyon-Tafani, Yousef Maali, Daniel Bouvard, Alan Diot, Frederic Laurent, Jerome Josse

## Abstract

*Staphylococcus aureus,* considered as a major human pathogen, is associated with severe infections such as bacteremia, endocarditis, skin and soft tissue infections, and bone and joint infections. Virulence mechanisms, such as biofilm formation and invasion/internalization of/into host cells, support the pathogenicity of *S. aureus* as they enable it to evade from the immune system and most antibiotic treatments. *S. aureus* can be internalized into non-professional phagocytic cells like fibroblasts, epithelial cells, endothelial cells and osteoblasts. The main pathway of internalization of *S. aureus* is FnBP-fibronectin-α5β1 integrin dependent. Interestingly, *S. pseudintermedius*, *S. delphini* and *S. argenteus* are able to invade osteoblasts, depending on the presence of FnBP-like proteins such as *Staphylococcus pseudintermedius* surface proteins D and L (SpsD/L) or *Staphylococcus delphini* surface protein Y (SdsY). However, the internalization capacity and mechanism have been poorly investigated in other staphylococci species. Here, we investigated the internalization capacity of staphylococci into osteoblasts at the genus level and attempted to correlate it with the presence of FnBP-like proteins by combining fibronectin adhesion assays, infection of osteoblasts and genome analysis. Over the 53 *Staphylococcus* species tested, half of them exhibited high internalization into osteoblasts. We highlighted that the “FnBP-fibronectin-α5β1 integrin” dependent internalization pathway of *S. aureus*, is well-conserved in 27 *Staphylococcus* species. *In silico* analysis identified multiple FnBP-like proteins correlating with the highly internalized species and showing diversity in their sequence organization, likely due to multiple acquisitions of such encoding genes throughout *Staphylococcus* evolution.

**AUTHOR SUMMARY:** *Staphylococcus aureus* is a pathogenic bacterium that causes severe infections including bone and joint infections. It invades bone cells, such as osteoblasts, using the bacterial surface protein FnBP, which binds to fibronectin, an extracellular compound, which subsequently binds to α5β1 integrin on the surface of osteoblasts. This cross-linking enables active internalization of *S. aureus* by the cells and potential intracellular persistence, which are responsible for the ability to induce staphylococcal chronic infections. While four species have been studied for their internalization into osteoblasts, a comprehensive genus-level investigation remains unexplored. Therefore, we investigated the conservation of this internalization capacity among the genus *Staphylococcus*, including a significant number of species of animal origin. Approximately half of the genus is capable of invading osteoblasts at varying levels via α5β1 integrin. Additionally, homologous proteins to FnBP were identified in most highly internalized species, suggesting a similar pathway of cell internalization to that of *S. aureus*. Genomic analysis reveals these proteins were acquired multiple times during evolution, suggesting they provide an advantage for host infection. In the context of One Health approach and the increasing number of animal pathogens causing human infections, understanding staphylococcal pathogenicity will help anticipate the emergence of new infectious diseases.

## INTRODUCTION

The genus *Staphylococcus* currently contains 67 validly published species or subspecies, most of which have been isolated from either human or animal samples [1,2]. Most of these species are commensal of the normal skin and mucosal microbiota. As opportunistic pathogens, they are involved in a wide range of human and animal infections [3,4]. Among them, *Staphylococcus aureus,* considered as the most frequent human opportunistic pathogen causing community and nosocomial infections, is associated with serious infections such as skin and soft tissue infections, ventilator-associated pneumonia (VAP), bacteremia, endocarditis, or bone and joint infections (BJI) [5]. However, the other staphylococcal species are also capable of causing human infections such as *Staphylococcus epidermidis* or *Staphylococcus capitis* involved in BJIs, *Staphylococcus lugdunensis* in pacemaker-associated infections, *Staphylococcus saprophyticus* in urinary tract infections, or even *Staphylococcus pseudintermedius*, initially isolated in animals, which can also cause human infections [6–9]. Virulence mechanisms, such as biofilm formation and invasion/internalization of/into host cells, have been proposed to promote staphylococcal infections, especially chronic forms, by allowing them to evade the immune system and most antibiotic treatments [10,11].

It is now well established that *S. aureus* has the ability to be internalized into non-professional phagocytic cells (NPPCs) such as fibroblasts, epithelial cells, endothelial cells and osteoblasts [12–15]. The main internalization pathway is related to the presence of the fibronectin binding proteins (FnBPA and FnBPB), adhesins located on the cell wall surface of the bacterium. This FnBP binds to fibronectin and subsequently to the cellular α5β1 integrin, thus acting as a bridge between the bacterium and the host cell by [16,17]. This specific interaction leads to the remodeling of the cytoskeleton, inducing cell membrane invagination. This allows *S. aureus* to enter host cells by endocytosis [18]. The two FnBPA and FnBPB isoforms of *S. aureus*, respectively encoded by the *fnbA* and *fnbB* genes [19,20], are part of the microbial surface components recognizing adhesive matrix molecules (MSCRAMM) family [21]. All the MSCRAMM proteins are composed of an N-terminal A domain with three subdomains N1, N2 and N3, the two last contributing to adhesion to fibrinogen, fibronectin-binding repeats located next to the A domain [22], a C-terminal Leu-Pro-X-Thr-Gly (LPXTG) motif enabling, with the signal sequence, the protein to be secreted and then covalently attached to peptidoglycans after sortase A cleavage [23]. In contrast, only limited data are available regarding the ability of the other staphylococcal species to be internalized into NPPCs. *S. lugdunensis* is able to invade epithelial and endothelial cells but not osteoblasts [24–26]. The internalization ability of *S. epidermidis* is more controversial. If Khalil *et al.* highlighted the invasion of osteoblasts but without the involvement of the α5β1 integrin [27], Valour *et al.* and Campoccia *et al.* reported that the invasion of osteoblasts by *S. epidermidis* is significantly lower compared to *S. aureus*, suggesting that this mechanism may not be the main pathogenic factor in orthopedic infections caused by *S. epidermidis* [26,28]. However, it has been clearly demonstrated that *S. pseudintermedius*, *Staphylococcus delphini* and *Staphylococcus argenteus* are able to invade osteoblasts [29–31]. Their internalization requires the presence of FnBP-like proteins such as surface proteins D and L (SpsD/L) in *S. pseudintermedius* or SdsY in *Staphylococcus delphini* [29–32]. To date, a global comprehensive investigation at the genus *Staphylococcus* level has not yet been carried out.

Therefore, we propose to investigate the capacity of internalization into osteoblasts of a large panel of species belonging to the genus *Staphylococcus* and attempted to correlate it with the presence of FnBP-like proteins. By combining genomic analysis, phenotypic fibronectin adhesion assays, and quantification of osteoblast invasion ability, we investigated the internalization pathway into osteoblasts among staphylococci. The 53 validly published staphylococcal species until 2020 have been involved, each species being represented by its reference strain. As a result, we have mapped the internalization capacity within the genus *Staphylococcus*. It appears that more than half of the species are capable of being internalized into osteoblasts, mainly through the “FnBP-like-fibronectin-α5β1 integrin”-dependent pathway. In addition, genomic data suggests that the FnBP-like proteins have been acquired on multiple occasions during *Staphylococcus* evolution.

## RESULTS

### *In silico* analysis of the staphylococcal reference strains

A neighbor-joining tree based on Mash genome distances was constructed using three *Salinicoccus* strains used as outgroup to root the tree. According to this mashtree (Fig 1), and on the contrary to Lamers *et al.* who defined 15 clusters [33], the genus *Staphylococcus* is divided into 7 clusters : extended *S. intermedius* cluster, *S. simulans*-*S. massiliensis* cluster, *S. auricularis* cluster, *S. edaphicus* cluster, *S. haemolyticus* cluster, *S. aureus* cluster, and *S. epidermidis* cluster. In particular, the *S. aureus* cluster comprises *S. aureus*, *S. argenteus*, *S. schweitzeri*, and *S. simiae*. The extended *S. intermedius* cluster contains notably *S. delphini* and *S. pseudintermedius*.

**Figure 1:**
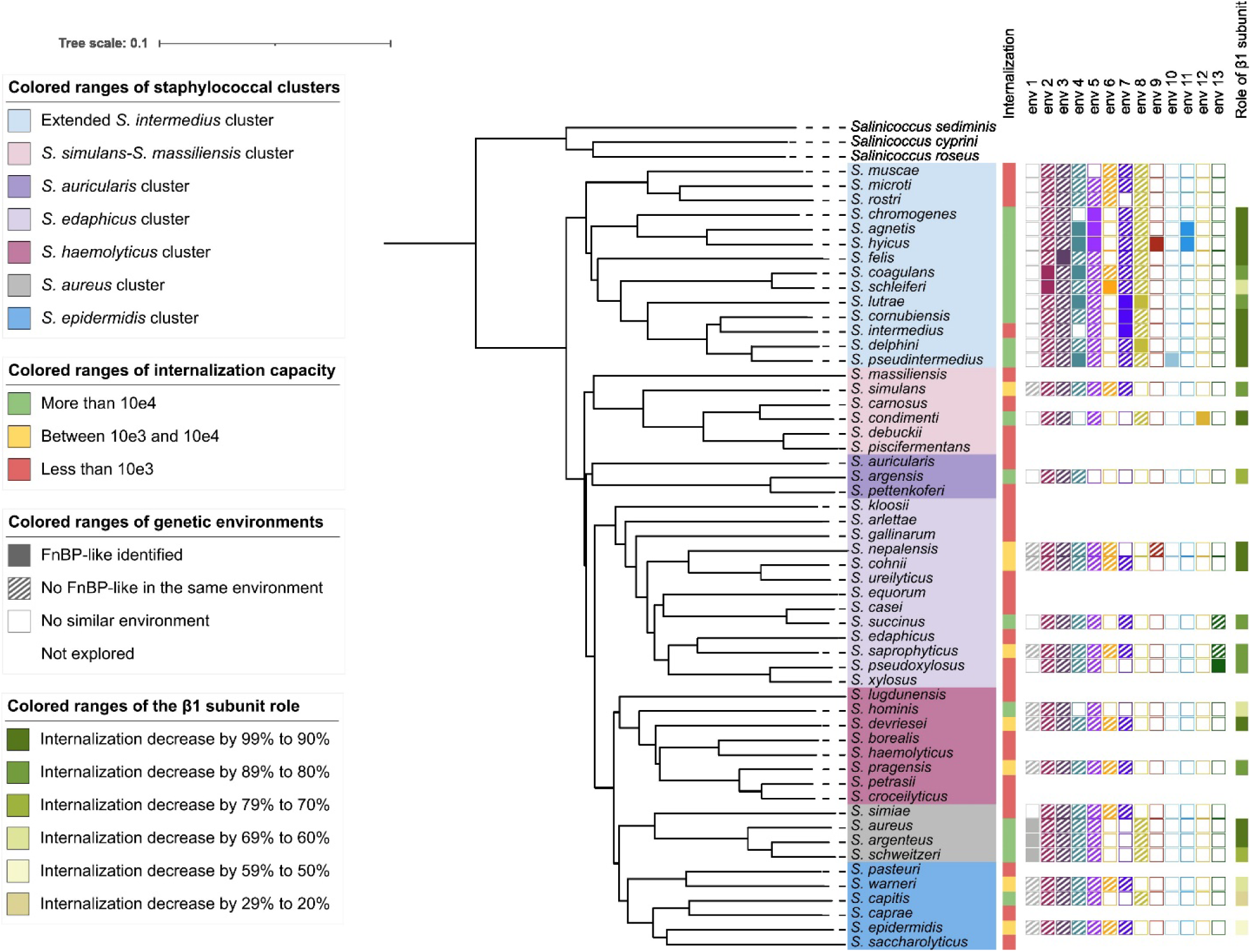
Neighbor-joining tree of the genus *Staphylococcus* and identification of FnBP homologous proteins. The tree based on Mash genome distances was constructed using mashtree v1.2.0 with default parameters. The internalization capacity, the presence of FnBP homologous proteins and their genetic environment, and the role of the β1 subunit of the α5β1 integrin in the internalization pathway of each species are illustrated. The internalization capacity of staphylococci has been evaluated using MG-63 human osteoblasts as previously described [30]. The “genetic environments” are composed of 10 genes: 5 upstream and 5 downstream of the FnBP homologous protein. An environment is considered not to be similar if genes from the reference environment are not found, or if identified but not in the same order or not organized together in the genome. The impact of the deletion of the β1 subunit of the α5β1 integrin is measured by the decrease in the proportion of staphylococcal internalization in murine osteoblasts lacking the β1 subunit (OB-β1^-/-^) compared to the wild-type murine osteoblasts (OB-β1^+/+^).

### Capacity to bind to human fibronectin and internalization into human osteoblasts

As the main pathway of internalization of *S. aureus* into osteoblasts is FnBP-fibronectin-α5β1 integrin dependent, adhesion to human fibronectin was quantified *in vitro* (Fig 2A). Thus, the residual adhesion capacity of the isogenic strain *S. aureus* DU5883, mutated for the two *fnb* genes of the positive control strain *S. aureus* 8325-4 [34], is reduced to 24.4% compared to the WT and was used as negative control. All reference strains with a higher fibronectin adhesion capacity than the negative control were considered to adhere to fibronectin. On the one hand, none of the strains from the *S. auricularis*, *S. epidermidis* and *S. haemolyticus* clusters adhere to the fibronectin *in vitro* under the experimental conditions, with adhesion rates not exceeding 19.7% compared to the positive control strain, except for the species *S. croceilyticus* belonging to *S. haemolyticus* cluster (47.5%). On the other hand, strains belonging to the *S. aureus*, *S. edaphicus* and extended *S. intermedius* clusters adhere moderately to strongly to fibronectin with rate of adhesion varying from 24.8% to 209.5% compared to the positive control strain. However, in some clusters such as the *S. simulans*-*S. massiliensis* cluster, some species adhere strongly, such as *S. condimenti* (206.8%), while others, such as *S. carnosus*, the closest clone based on phylogeny to *S. condimenti*, do not adhere to fibronectin at all (Fig 2A). These results highlighted that the *in vitro* fibronectin adhesion capacity levels are highly variable for both intra- and inter-cluster staphylococcal strains.

**Figure 2:**
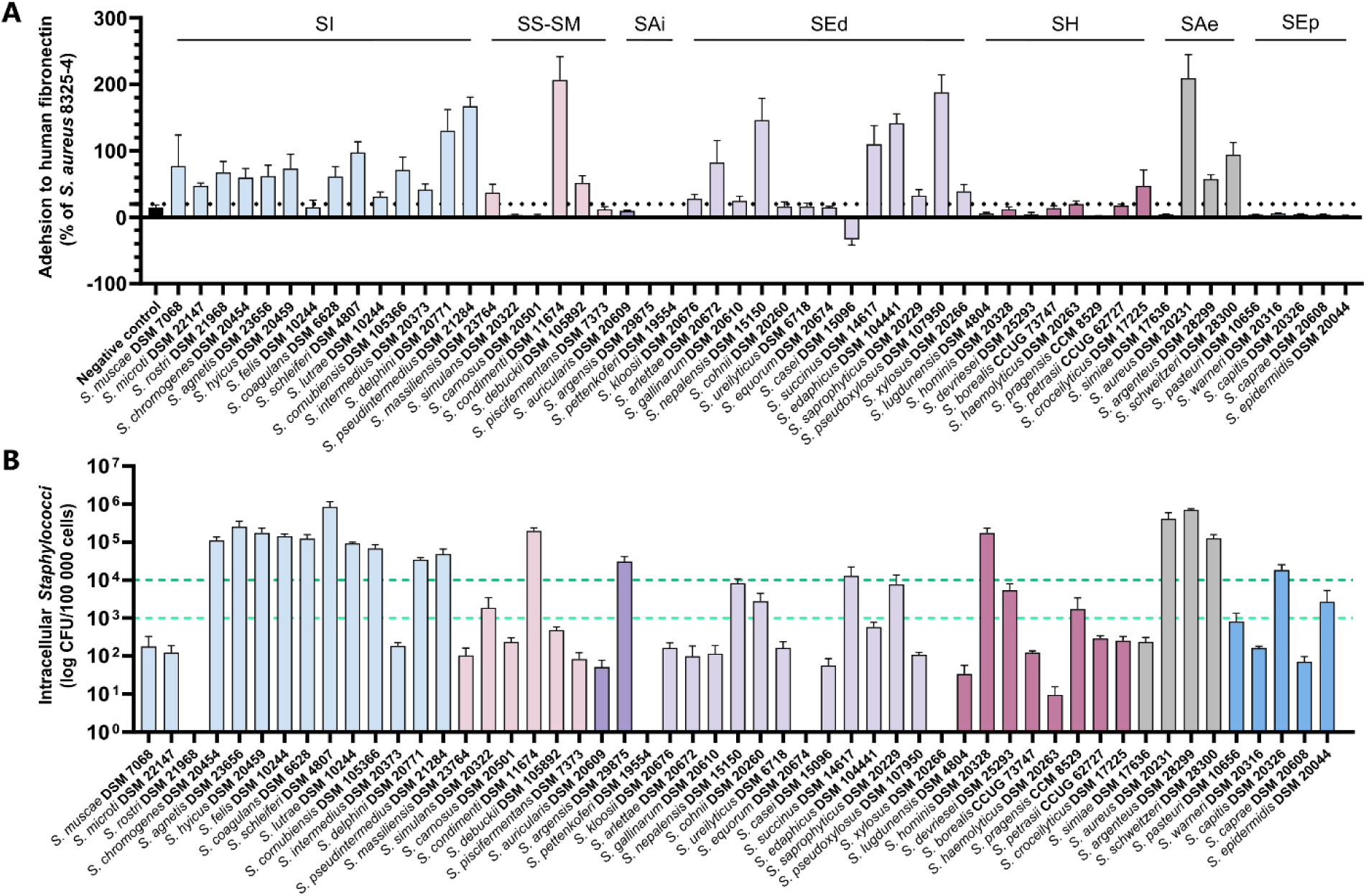
Assessment of human fibronectin adhesion and internalization into human osteoblasts for *Staphylococcus spp.* Experiments have been performed at least three times independently. Results are represented as the mean with the standard error of the mean. A: *In vitro* fibronectin adhesion capacity of 53 refence strains of staphylococci validly published until 2020. The data are expressed in percentage of the adhesion ability compared to the strain *S. aureus* 8325-4 (Positive control). *S. aureus* DU5883 strain, which is mutated on the two genes *fnbA* and *fnbB* coding for the two isoforms FnBPA and FnBPB was used as negative control. B: Quantification of staphylococci internalization into MG-63 human osteoblasts. The threshold to characterize low and high internalization capacity are respectively 10^3^ and 10^4^ intracellular staphylococci per 10^5^ infected cells. SI: extended *S. intermedius* cluster; SS-SM: *S. simulans*-*S. massiliensis* cluster; SAi: *S. auricularis* cluster; SEd: *S. edaphicus* cluster; SH: *S. haemolyticus* cluster; SAe: *S. aureus* cluster; SEp: *S. epidermidis* cluster.

As several *Staphylococcus* species have been shown to be able to be internalized into osteoblasts, the internalization capacity of all the reference strains was tested using MG-63 osteoblasts (Fig 2B). The thresholds for characterizing low and high internalization capacity were arbitrarily defined as <10^3^ and >10^4^ intracellular staphylococci per 10^5^ infected cells respectively, to facilitate interpretation based on our experience with staphylococcal internalization.

Within the *S. aureus* cluster, *S. aureus*, *S. argenteus* and *S. schweitzeri* were highly internalized, whereas *S. simiae* was weakly internalized into osteoblasts. Interestingly, *S. simiae* is on an independent branch of the phylogenetic tree compared to the three other species of this cluster (Fig 1). Together these data suggest that the ability to be internalized was acquired after the genetic drift inducing speciation of *S. simiae* within the *S. aureus* cluster.

Similarly, most species within the extended *S. intermedius* cluster exhibit a high capacity for internalization into osteoblasts. However, there were exceptions such as the species *S. muscae*, *S. microti,* and *S. rostri*, which exhibit weak or almost no internalization (Fig 2B). Again, these three species belonged to the same phylogenetic branch compared to the other species of the extended *S. intermedius* cluster (Fig 1, first column). Together, the species within these two clusters, the *S. aureus* cluster and the extended *S. intermedius* cluster, have a high internalization capacity reaching 8.5×10^5^ intracellular staphylococci per 10^5^ infected cells for *S. schleiferi*. The case of *S. intermedius* which was weakly internalized, is discussed in more details below (see section Reduced internalization due to truncated FnBP-like protein).

In the other clusters, most of the reference strains are not able to be internalized. However, *S. condimenti* in the *S. simulans*-*S. massiliensis* cluster, *S. argensis* in the *S. auricularis* cluster, *S. succinus* in the *S. edaphicus* cluster, *S. hominis* in the *S. haemolyticus* cluster, and *S. capitis* in the *S. epidermidis* cluster are all five highly capable of internalization into osteoblasts. Their internalization capacity was respectively of 2.0×10^5^, 3.1×10^4^, 1.3 ×10^4^, 1.7×10^5^, and 1.8 ×10^4^ intracellular staphylococci per 10^5^ infected cells (Fig 2B). These five highly internalized species are not present on a distinct branch within their respective clusters (Fig 1). This suggests that their internalization phenotype may have emerged spontaneously during the evolution of the species within their own genetic cluster, and sporadically in contrast to the extended *S. intermedius* and *S. aureus* clusters.

In addition, the species *S. simulans*, *S. nepalensis*, *S. cohnii*, *S. saprophyticus*, *S. devriesei*, *S. pragensis*, and *S. epidermidis* have an average internalization capacity varying from 1.7×10^3^ to 8.4×10^3^ intracellular staphylococci per 10^5^ infected cells, e.g between the two thresholds arbitrarily chosen to discriminate high and low internalization rate (Fig 2B). All these species are present in distinct clusters. Like *S. condimenti*, *S. argensis*, *S. succinus*, *S. hominis*, and *S. capitis*, these species are not located on a distant/independent branch within their respective clusters (Fig 1). In the same way, it suggests that the internalization phenotype may have emerged spontaneously and sporadically within their respective clusters. Of note, some species are not able at all to be internalized into MG-63 osteoblasts at all regardless of the cluster group, namely the *S. pettenkoferi*, *S. equorum*, *S. xylosus* and *S. rostri* (Fig 2B).

The ability to adhere to fibronectin and internalize into osteoblasts are both highly variable from one species to another and no direct correlation between these two characteristics was observed. For instance, all the strains from the extended *S. intermedius* cluster showed a high level of internalization into osteoblasts except *S. muscae*, *S. microti S. rostri*, and *S. intermedius*. Interestingly, *S. felis* poorly adheres to fibronectin (only 15.0% compared to *S. aureus* 8325-4 positive control strain) but has an internalization capacity greater than 10^5^ intracellular staphylococci per 10^5^ infected cells. Likewise, in the *S. aureus* cluster, *S. argenteus* is the species with the highest level of internalization, but not the species with the highest level of adhesion to fibronectin (Fig 2).

### *In silico* identification of *fnb* gene homologues and analysis of their genomic environment

*In silico* analyses have been carried out to determine whether the internalization of species (n=27) could be explained by the presence of FnBP homologs. The analysis of sequence homology between FnBPA of *S. aureus* and protein sequences from genomes of *Staphylococcus* strains with an internalization level higher than 10^3^ intracellular staphylococci per 10^5^ infected cells identified 33 candidates. FnBPA is characterized by the presence of 4 main domains: YSIRK Gram-positive signal peptide, SDR-like Ig domain, fibrinogen-binding domain 2 and LPXTG cell wall anchor domain. Three candidates lacking these domains and four presenting serine aspartate (SD) repeats characteristic of serine-aspartate repeats protein Sdr, were removed.

Then, the genetic environment of the 26 remaining candidate genes were determined to check if some of these genes were located in similar environments (S1 Table). A “genetic environment” was defined by the 5 genes upstream and 5 genes downstream of the FnBP-like protein. An environment was considered non-similar if no genes from the reference environment are found, or if the identified genes are not in the same order or are not organized together in the genome. The systematic search for each of these “genetic environments” in all staphylococci genomes and if present, the examination of their gene contents, allowed us to identify additional potential candidates that were not previously identified based on sequence homology analysis. Thus, a further set of 6 additional genes were identified by this method (i.e., genes in the same position within the same genetic environment as FnBP-like identified by protein homology in other *Staphylococcus* species), but 3 of these were removed because they did not contain the specific-FnBP protein domains. In total, 29 potential FnBP-like proteins were identified. All proteins were named with “S” for *Staphylococcus*, two letters from the species name, and “s” for surface protein (S2 Table).

The protein domains of the FnBP-like proteins were highly similar (S1 Fig). All proteins contain a signal peptide, an A domain consisting of the domains N1, N2 and N3, a repeat-containing domain, and an LPXTG motif, except ScasB of *S. coagulans*, which has an LPXSG motif. Some proteins, namely ScasB, SslsA, SlusB, ScrsA, SinsA, SdesA, and SpisA, present in *S. coagulans*, *S. schleiferi*, *S. lutrae*, *S. cornubiensis*, *S. intermedius*, *S. delphini*, and *S. pseudintermedius* respectively, contain several copies of the SdrB domain in their repeated region, which is characteristic of the Clf-Sdr family proteins and collagen adhesin Cna [35]. All the FnBP-like proteins contain in their C-terminal region a proline-rich repeat domain (P), a wall-spanning domain (W), and a membrane-spanning domain (M), except SagsC from *S. agnetis*, SlusA from *S. lutrae*, and ScnsA from *S. condimenti*.

In the *S. aureus* cluster, the two FnBP-like proteins (FnBPA and FnBPB-like proteins) are identified in a single similar environment in *S. aureus*, *S. argenteus*, and *S. schweitzeri* species (Fig 1, S1 Table), whereas in *S. simiae*, the only species unable to internalize into osteoblasts, the same genetic environment is present but lacks any genes encoding FnBP-like proteins.

On the contrary, in the extended *S. intermedius* cluster, FnBP-like proteins have been identified in several different environments. Interestingly, *S. schleiferi* and *S. coagulans* share environment 2, these species being previously described as two subspecies *S. schleiferi* subsp. *schleiferi* and *S. schleiferi* subsp. *coagulans* [1]. Environment 5 is common to *S. chromogenes*, *S. agnetis*, and *S. hyicus*, which are present in the same root on the neighbor-joining tree but harbored three different FnBP-like proteins (Fig 1). Environments 4 and 11 are also common to *S. agnetis* and *S. hyicus*, which are phylogenetically close. Conversely, environment 7 is similar in *S. lutrae*, *S. cornubiensis*, and *S. intermedius*, while these species do not share a close common ancestor. Environment 3 is present in all investigated species, but it harbors a FnBP-like protein only in *S. felis*. Surprisingly, an FnBP-like protein has been identified in *S. condimenti* in environment 12, only retrieved in this species. Even more surprising, an FnBP-like protein has been identified in *S. pseudoxylosus*, whereas this species is not able to be internalized into osteoblasts. However, the level of adhesion to fibronectin of *S. pseudoxylosus* is well above the *S. aureus* 8325-4 control strain, suggesting that this FnBP-like protein could contribute to its adhesion to fibronectin, but our data indicates that it is defective to contributing to internalization into osteoblasts. Nevertheless, no FnBP-like proteins have been identified for *S. argensis*, *S. succinus*, *S. hominis*, and *S. capitis*, while they demonstrate a high capacity of internalization ranging from 1.2×10^4^ to 1.7×10^5^ intracellular staphylococci per 10^5^ infected cells (Fig 2B). In addition, no FnBP-like proteins have been identified for the strains presenting an average capacity of internalization (10^3^-10^4^ intracellular staphylococci per 10^5^ infected cells) into human osteoblasts either (Fig 1).

To better understand the evolutionary relationship within FnBP-like proteins, two molecular phylogenetic trees have been constructed: one based on the entire protein sequences (part A in S2 Fig), the other focused on the A domains (part B in S2 Fig). The distributions are largely overlapping, with genetic environments similarly distributed between the two trees. In the evolutionary tree based on the A domain (S2 Fig A), the FnBP-like proteins identified within the *S. aureus* cluster are grouped on a branch subdivided into two branches, one branch including isoforms A and the other including isoforms B, but all were inserted in a same common genetic environment. The A domains of the FnBP-like proteins identified in the extended *S. intermedius* cluster are distributed in the two main branches, the upper branch containing only proteins whose repeated region contains SdrB domains (S1 Fig). We noticed that the FnBP-like proteins inserted in the genetic environment 4 are divided, with 3 on the upper branch and 2 on the lower branch, while on the lower branch, all proteins are grouped according to the genetic environment in which they were identified. Additionally, the lower branch contains proteins from the *S. aureus* cluster, encompassing the two isoforms of the FnBP that were initially identified. This suggests that the proteins on the lower branch belong to the FnBP family, while the upper branch contains mostly Sdr or Cna type proteins due to the presence of SdrB domains. Surprisingly, the tree representing the evolution of domain A (S2 Fig A) is highly similar to the tree representing the evolution of the entire protein sequences (S2 Fig B). The repeated region being particularly variable, the sequence alignment contains many gaps. Therefore, it is likely that the sequences shared by the 29 proteins, which form the basis of the complete protein tree, are highly similar to the A domain sequences. Lastly, it is noteworthy that the proteins are not distributed according to the same way as in the species phylogenetic tree and that they are predominantly clustered according to the genetic environments in which they were discovered. This supports the hypothesis of multiple acquisitions independently of the evolution of the *Staphylococcus* species and genus.

Intriguingly, each of the genetic environments harboring FnBP-like protein contains in addition at least one other virulence factor, regulator, or metalloprotease gene (S1 Table). These virulence factors are known to be involved in either biofilm formation, toxin secretion, natural competence, or stress response. Besides, environments 1, 2, 3, and 4 contain the three categories of genes. This suggests that FnBP-like proteins might be located in pathogenicity islands.

### Characterization of the role of the β1 subunit in the internalization pathway into osteoblasts

To be internalized into osteoblasts, *S. aureus* requires the presence of α5β1 integrin on the surface of the host cells. Thus, to further characterize the internalization pathway into osteoblasts of staphylococci, internalization assays has been performed with murine osteoblasts deleted or not for the β1 subunit of the α5β1 integrin (respectively, OB-β1^-/-^ and OB-β1^+/+^) as previously described [30], the internalization capacity of staphylococci being supposed to be reduced in the absence of the β1 subunit of the α5β1 integrin. We first investigated the impact of the β1 subunit of the α5β1 integrin in the internalization mechanism for species showing an internalization capacity higher than 10^4^ intracellular staphylococci per 10^5^ infected cells (Fig 3A). The internalization capacity of *S. aureus* strain, used as control, decreases by 96.6% in osteoblasts OB-β1^-/-^ deleted for the β1 subunit compared to wild type osteoblasts OB-β1^+/+^. The number of internalized staphylococci in osteoblasts OB-β1^-/-^ significantly decreases compared to the internalized staphylococci in osteoblasts OB-β1^+/+^ for 16 *Staphylococcus* species, with a loss of internalization capacity ranging from 64.1% for *S. hominis* to 98.1% for *S. chromogenes*. However, the internalization capacities of *S. schleiferi* and *S. capitis* are not significantly impacted by the absence of the β1 subunit of the α5β1 integrin. Of note, a FnBP-like protein has been identified in the genome of all these species, except for *S. capitis*. This result suggests that the internalization pathway of all these species is similar to the one of *S. aureus* and involved the interaction of a FnBP-like protein and the α5β1 integrin, except for *S. capitis*.

**Figure 3:**
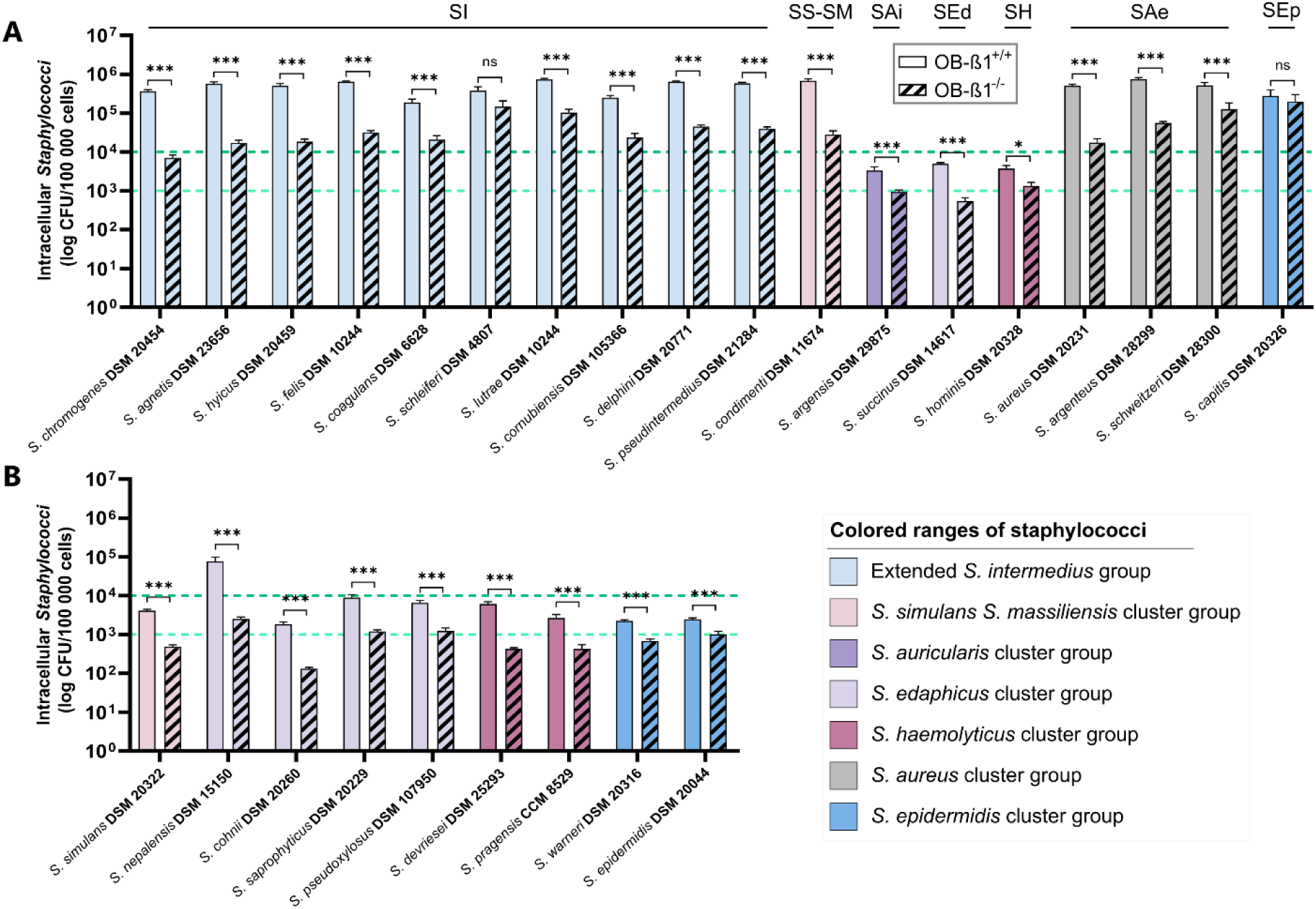
Characterization of the role of the β1 subunit of the α5β1 integrin in the internalization pathway. The role of the β1 subunit of the α5β1 integrin was investigated using OB-β1^-/-^ deleted for the β1 subunit of the α5β1 integrin and murine wild type osteoblasts OB-β1^+/+^. The experiments have been performed at least three times independently. Results are represented as the mean with the standard error of the mean. A: Characterization of the role of the β1 subunit of the α5β1 integrin in the internalization pathway of staphylococci with high internalization capacity in MG-63 human osteoblasts. B: Characterization of the role of the β1 subunit of the α5β1 integrin in the internalization pathway of staphylococci with average internalization capacity in MG-63 human osteoblasts. Mann-Whitney test: p-value non-significant (ns), 0.01 (*), 0.001 (**), 0.0001 (***). SI: extended *S. intermedius* cluster; SS-SM: *S. simulans*-*S. massiliensis* cluster; SAi: *S. auricularis* cluster; SEd: *S. edaphicus* cluster; SH: *S. haemolyticus* cluster; SAe: *S. aureus* cluster; SEp: *S. epidermidis* cluster.

The role of the β1 subunit of the α5β1 integrin in the internalization mechanism of the reference strains with an average internalization between 10^3^ and 10^4^ intracellular staphylococci per 10^5^ infected cells has also been investigated. The internalization pathway of two additional species (*S. pseudoxylosus* and *S. warneri*), with an internalization capacity close to the minimum threshold of 10^3^ intracellular staphylococci per 10^5^ infected cells, was explored too (Fig 3B). The proportion of internalized staphylococci also significatively decreases for the 9 reference strains tested, ranging from 59.0% for *S. epidermidis* to 96.7% for *S. nepalensis*. No FnBP-like protein has been identified in the genome of these reference strains, except for *S. pseudoxylosus*. Of note, the repeated region of SpxA does not seem to contain Fn-binding repeats according to a domain search performed on InterPro website (S1 Fig). The internalization pathway of the strains requires the cell receptor α5β1 integrin, but another bacterial adhesin could be involved.

### Reduced internalization due to truncated FnBP-like protein

The reference strain *S. intermedius* DSM 20373 presented an atypical behavior within the extended *S. intermedius* cluster as it is not able to be internalized into human osteoblasts despite the presence of FnBP-like gene in its genome. The fibronectin adhesion assays of 8 *S. intermedius* strains showed a high variability within strains with proportion of adherent bacteria compared to the positive control *S. aureus* 8325-4 strain ranged from 8.91% (e.g. below the negative control strain *S. aureus* DU5883) to 84.9% (Fig 4A). The internalization capacity varied from 1.5×10^1^ to 3.5×10^3^ intracellular staphylococci per 10^5^ infected cells (Fig 4B). There was no correlation between adhesion to fibronectin and internalization into osteoblasts. Strains that adhere the least were not necessarily the ones that internalize the least and *vice versa*. Finally, for *S. intermedius* DSM 20373, the proportion of internalized bacteria decreased by 92.8% suggesting that the residual internalization of bacteria is also dependent on the α5β1 integrin.

**Figure 4:**
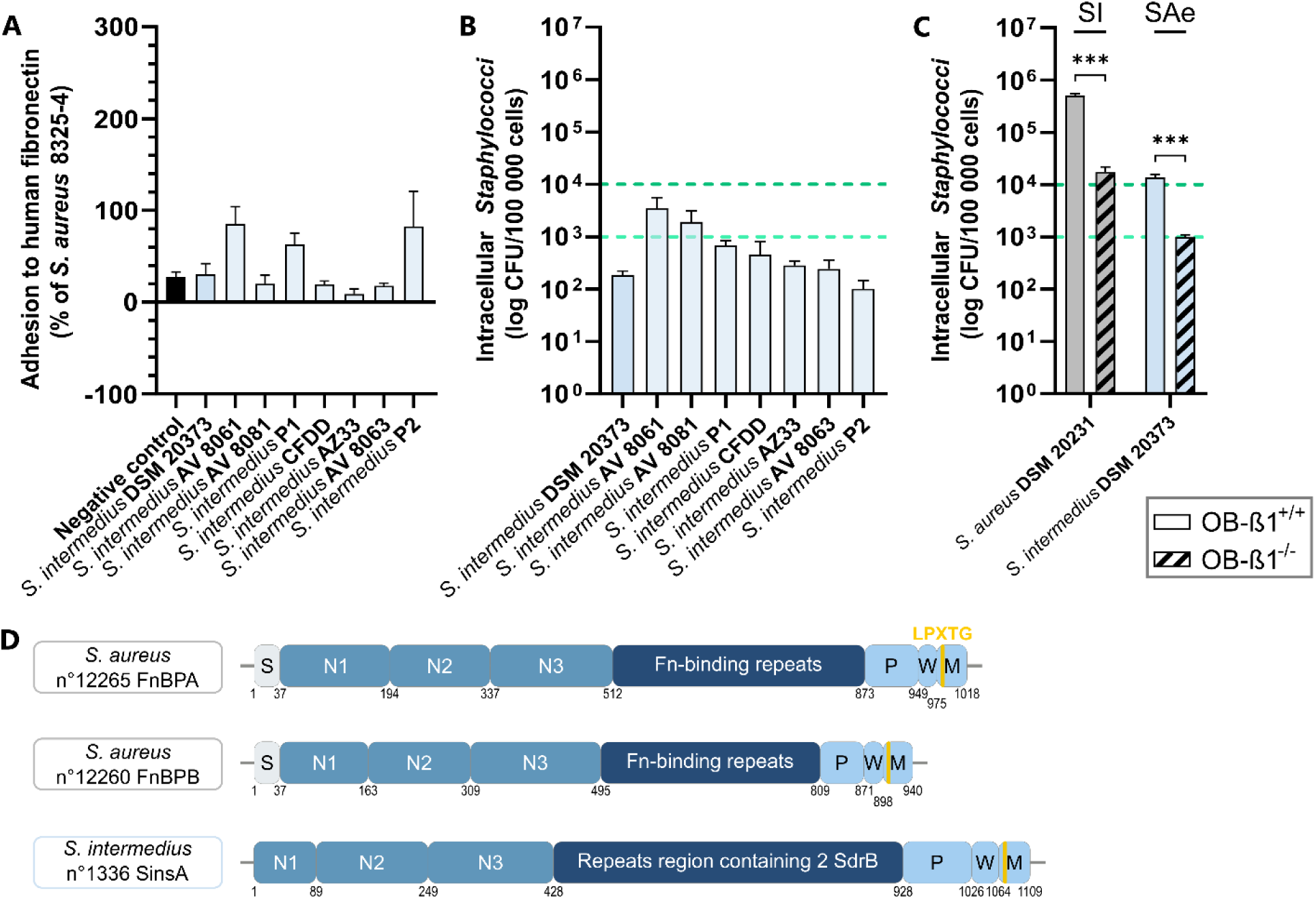
Characterization of the internalization pathway of *Staphylococcus intermedius*. The experiments have been performed at least three times independently. Results are represented as the mean with the standard error of the mean. A: Evaluation of adhesion to fibronectin of *S. intermedius*. The data are expressed in percentage of the adhesion ability of the positive control strain *S. aureus* 8325-4. The negative control strain corresponds to the fibronectin adhesion capacity of the *S. aureus* DU5883 strain, which is deleted of the two genes *fnbA* and *fnbB* coding for the two isoforms FnBPA and FnBPB. B: Quantification of internalization into MG-63 human osteoblasts of *S. intermedius*. C: Characterization of the role of the β1 subunit of the α5β1 integrin in the internalization pathway of *S. intermedius*. The role of the β1 subunit of the α5β1 integrin was investigated using OB-β1^-/-^ deleted for the β1 subunit of the α5β1 integrin and murine wild type osteoblasts OB-β1^+/+^. SI: extended *S. intermedius* cluster; SAe: *S. aureus* cluster. Mann-Whitney test: p-value non-significant (ns), 0.01 (*), 0.001 (**), 0.0001 (***). D: Domain organization of homologous protein of *S. intermedius* of FnBP. S: signal peptide; N1, N2 and N3 represent the A domain; Fn-binding repeats: fibronectin binding protein; P: proline-rich repeats; W: wall-spanning domain; M: membrane-spanning domain. The protein numbers correspond to the sequences that were identified in the *in silico* analysis.

*In silico* analysis of *S. intermedius* DSM 20373 genome identified a homologous protein to FnBP, which we named *Staphylococcus intermedius* surface protein SinsA (Fig 4D). This protein is composed of the A domain made of three subdomains N1, N2 and N3, a repeated region that includes two SdrB domains, and an LPXTG motif in its C-terminal region. On the contrary to *S. aureus* FnBPA/B, SinsA lacks a signal peptide sequence. The absence of signal peptide and the presence of SdrB domains in the repeated region could influence the internalization capacity of *S. intermedius* into osteoblasts.

## DISCUSSION

Bacterial internalization is a virulence mechanism shared by several genus and species, such as *Legionella pneumophila*, *Listeria monocytogenes* or *S. aureus* [36–39]. In particular, *S. aureus* internalization into human NPPC, and notably into osteoblasts, has been extensively explored [11], on the contrary to other *Staphylococcus* species. Internalization mechanism is particularly of concern since persistence into the intracellular niche has been reported to be involved in the physiopathology of chronic staphylococcal infections [40]. Indeed, intracellular localization enables bacteria to be protected from the immune system of the infected host and from the activity of most antibiotics [10,40]. Furthermore, if *S. epidermidis, S. lugdunensis, S. capitis* are well known to induce human infections, many other non-*aureus* staphylococci such as *S. condimenti*, *S. cohnii* are increasingly reported [41–51], notably in BJIs but also pneumonia, endocarditis or endophthalmitis [45,50,51].

In this context, it seemed relevant to us to evaluate the internalization capacity of staphylococci in NPPCs on a genus scale. We selected osteoblasts as host cells of interest given that staphylococci are frequently implicated in BJIs.

Globally, our findings demonstrate that the internalization of staphylococci into osteoblasts is species-dependent and that the internalization pathway initially described for *S. aureus*, the FnBP-fibronectin-α5β1 integrin, is also used by most species capable of being internalized into osteoblasts. Furthermore, most species capable of being internalized belong to two distinct genetic clusters, the *S. aureus* cluster and the extended *S. intermedius* (Fig 1).

The *S. aureus* cluster can be considered as the textbook case of our study. This cluster is composed of 4 species: *S. aureus*, *S. argenteus*, *S. schweitzeri* and *S. simiae*. *S. aureus*, *S. argenteus* and *S. schweitzeri* are all three capable of being internalized by osteoblasts unlike *S. simiae*. *S. aureus*, *S. argenteus* and *S. schweitzeri* all have two FnBP or FnBP-like isoforms. The genes coding for these two isoforms are contiguous and inserted in a similar genetic environment for the three species. On the contrary, *S. simiae* does not present any gene encoding a FnBP-like within its genome. *S. simiae* has the genetic environment in which FnBP or FnBP-like have been identified in *S. aureus*, *S. argenteus* and *S. schweitzeri* but without the genes encoding the latter. On the scale of the phylogeny, *S. simiae* is on a distinct branch compared to the other 3 species. Taken together, these observations support the hypothesis that internalization capacity is a trait acquired by a common ancestor of *S. aureus*, *S. argenteus* and *S. schweitzeri* after their phylogenetic divergence from *S. simiae*.

This hypothesis agrees with a previous comparative genomics study by Suzuki *et al.* which suggests that *S. aureus* has acquired virulence factors through horizontal gene transfer, after the split of *S. aureus* and *S. simiae* from their common ancestor [52]. Note that currently no published study has focused on the internalization of *S. simiae*, whatever the host cell type.

Regarding *S. schweitzeri*, our results align with those of Grossmann *et al.* who reported an internalization capacity in Vero cells like that of *S. aureus* for 58 strains of *S. schweitzeri.* As raised by a recent review, the internalization capacity of *S. schweitzeri* comparable to that of *S. aureus* questions the possibility of this species to become an emerging zoonotic pathogen of interest [53].

The second cluster highlighted by our study is the extended *S. intermedius* cluster. It is composed of 14 species, mostly initially isolated in animal samples [4] Among these 14 species, 10 showed significant internalization capacity within osteoblasts (Fig 1). Regarding species that cannot be internalized, no FnBP-like protein has been identified for *S. muscae*, *S. microti* and *S. rostri*. As observed previously for *S. simiae* in the *S. aureus* cluster, these 3 species are grouped together on a branch which separates early from the rest of the cluster. Again, in this case, the hypothesis would be that the genes responsible for internalization into osteoblasts were acquired by the other staphylococci of this cluster after the phylogenic divergence with *S. muscae*, *S. microti* and *S. rostri*.

However, for this cluster the case is more complex than for the *S. aureus* cluster. Indeed, the FnBP-like identified for the 10 strains capable of being internalized were identified in different genetic environments. Some FnBP-like proteins are common between genetically close species, such as SchsA, SagsB and ShysB which are respectively the FnBP-like of *S. chromogenes*, *S. agnetis* and *S. hyicus* identified in the same genetic environment 5. Additionally, according to the phylogenetic tree of FnBP-like protein, proteins in the environments 2, 3, 4, 9, and 11 are quite similar (S2 Fig), which is corroborated with the organization of functional domains (S1 Fig). This complex distribution of FnBP-like proteins across diverse genetic contexts, combined with their frequent association with antibiotic resistance genes, metal-related elements, and mobile genetic elements (S1 Table), supports the hypothesis of horizontal gene transfer. Such transfer events may have contributed to the dissemination of these proteins and shaped their current distribution among different genetic environments. Analyzing insertion sites in these genetic environments could provide valuable insights.

Besides, *S. intermedius* is not or weakly able to be internalized into osteoblasts depending on the strains, while an FnBP-like protein, SinsA, has been identified in the reference strain genome. As SinsA lacks signal peptide, we hypothesized that the absence of a signal peptide could explain the observed phenotype. However, given that the FnBP-like protein of *S. delphini* SdesA exhibits a similar case, we can no longer be certain about this assumption. Besides, it is noteworthy that the SdsY protein reported in Maali *et al.* was not identified in this new analysis [30].

Two other FnBP-like, Scns and SpxsA, were identified in *S. condimenti* and *S. pseudoxylosus* respectively, two species located outside the *S. aureus* and extended *S. intermedius* clusters. Within their respective clusters, these 2 species are the only ones to have an FnBP-like protein.

Concerning *S. condimenti*, this species exhibited a strong internalization capacity in osteoblasts. ScnsA, the FnBP-like identified in this species, is located in a genetic environment unique to it. Phylogenetic analysis specific to FnBP-like showed that this protein was genetically close to SchsA, SagsB and ShysB, FnBP-like proteins identified in *S. chromogenes*, *S. agnetis* and *S. hyicus* in genetic environment 5 (Fig 1, S2 Fig). It can be easily imagined that the FnBP from one of the 3 previously mentioned species was transferred horizontally to *S. condimenti* and was inserted into a different genetic environment.

A similar situation is observed for *S. pseudoxylosus*, except that this species is not internalized by osteoblasts. Interestingly, the FnBP-like identified for this species, SpxsA, is genetically close to ScnsA identified in *S. condimenti* and to SchsA, SagsB and ShysB identified in *S. chromogenes*, *S. agnetis* and *S. hyicus* respectively. The hypothesis of a horizontal gene transfer with insertion in a genetic environment different from that of origin can once again be raised. Concerning the absence of internalization in osteoblasts, the repeated region of this protein contains an atypical keratinocyte proline rich domain. As the number of repeats is correlated to fibronectin adhesion [54], the lack of fibronectin-binding repeats of the SpxsA protein might explain *S. pseudoxylosus* internalization disability.

Twelve other species demonstrated an average or strong internalization capacity within osteoblasts. For these species, no FnBP-like could be identified directly in the genomes or during in-depth analyses of the genetic environments identified in the other species. These results are consistent with the absence of adhesion to fibronectin observed for these species, except for *S. cohnii*. However, on the osteoblast side, the internalization of these species seems to be generally dependent on the presence of the α5β1 integrin. Some authors have already proposed that other proteins able to bind to α5β1 integrin are involved in the internalization pathway of non-*aureus* staphylococci such as Atl, which mediates the *S. epidermidis* uptake into endothelial cells via the cell receptors α5β1 integrin or Hsc70 [55].

Our study is not the first to look at a virulence factor at the scale of the genus *Staphylococcus*. Pickering *et al*., in their 2021 study, focused on the distribution of plasma clotting capacity associated with the *vwb* gene coding for the von Willebrand factor-binding protein (vWbp) [56]. Interestingly, all the strains identified as able to clot plasma in their study are also those highly able to be internalized into osteoblasts in our study, except for the particular case of *S. intermedius*. The authors suggested that this phenotype could have appeared through species evolution to adapt to its host [56]. In the same way, FnBP-like proteins are identified in several clusters within the genus *Staphylococcus* and correspond to species able to be internalized into osteoblasts. Like vWbp, FnBP-like proteins could be a key virulence factor in host-pathogen interactions within the genus *Staphylococcus*.

Several limitations must be considered to have a critical vision of our study. The major limitation of our work is that we only studied the reference strain of each species. The generalization of our observations to all strains of a species should therefore be taken with caution. Indeed, it is recognized that the ability of *S. aureus* to invade host cells is strain-dependent, with widely varying internalization capacities depending on the strain studied [57,58]. This strain-dependent internalization is illustrated in our study by our data specific to *S. intermedius* species, whose internalization varies by more than one log depending on the strain studied. For future studies, it would be interesting to focus on certain species but to include a collection of strains for each of them in order to take into account intraspecies diversity.

We focused our work on internalization in osteoblasts and we used the MG-63 cell line to carry out our experiments. On the one hand, the use of primary osteoblasts could have allowed us to have a closer approach to the physiopathological reality. However, the culture of primary cells is more demanding than that of cell lines, so it would have been difficult to correctly carry out cell infections for 53 strains using primary osteoblasts. On the other hand, it would have been interesting to test the internalization of the 53 strains of staphylococci in another cellular model such as keratinocytes or endothelial cells, to question the host cell character dependence on our results. We can easily assume that the classification of different species based on their internalization capacity would have varied depending on the host cell type. Strobel *et al.* notably observed that the amount of internalized *S. aureus* varies depending on the cell types and whether or not experiments are done with primary cells, as well as cytotoxicity and persistence into host cells [57]. In our study, the variations in internalization for *S. intermedius* between OB-β1^+/+^ murine osteoblasts and MG-63 human osteoblasts illustrate this point well and even questions the role of the human or animal nature of the host cells.

The definition of genetic environments in the analysis of different FnBP-like proteins could also be considered as a limit of our work. These environments must be considered with caution as we arbitrarily define them by considering 10 genes on either side of the FnBP-like protein. Because other genes could have been acquired in combination with the FnBP-like protein, a possible shift in the original environment could happen, leading to a misinterpretation of a genetic environment as different.

A final major point, which goes beyond the scope of our study, is the classification of proteins identified as FnBP-like. Among the identified FnBP-like proteins, 8 proteins contain at least one SdrB domain. According to Foster, SdrB domain, also named B repeats or B domain, is common to Clf, Cna and Sdr proteins of the MSCRAMMs family [59]. In particular, Cna repeated region is composed of four B repeats, while Sdr and ClfA comprise two B repeats [22]. This suggests that the FnBP-like proteins ScasB, SslsB, SlusB, ScrsA, SinsA, SdesA, and SpisA identified in the *S. coagulans*, *S. schleiferi*, *S. lutrae*, *S. cornubiensis*, *S. intermedius*, *S. delphini*, and *S. pseudintermedius* strains could belongs to Clf, Cna or Sdr type. Interestingly, Josefsson *et al.* and Patti *et al.* identified Cna and ClfA as virulent determinants in the context of staphylococcal BJIs [60,61]. These observations raise interest in investigating the role of Cna and ClfA in an internalization pathway dependent on the α5β1 integrin.

However, as it is difficult to classify the identified proteins into one category or another, the current classification is likely a bit too strict. In-depth studies are required to more easily discriminate protein categories and/or to determine whether a new category should be defined.

In conclusion, the present study investigates bacterial internalization among the genus *Staphylococcus*, using osteoblasts as a model cell line. The internalization capacity varies across species. Over 53 *Staphylococcus* species tested, half of them exhibit high internalization into osteoblasts. The main internalization pathway of *S. aureus*, “FnBP-fibronectin-α5β1 integrin” dependent, is well-conserved in 27 *Staphylococcus* species. *In silico* analysis identified 29 FnBP-like proteins, showing diversity in their sequence organization, which could be due to multiple acquisitions and/or horizontal gene transfer between species throughout *Staphylococcus* evolution.

## MATERIALS AND METHODS

### Phylogenetic analysis

Genomes in fasta format were downloaded from NCBI (S3 Table). Genomes were annotated using Bakta v1.6.1 with default parameters. A neighbor-joining tree based on Mash genome distances was constructed with mashtree v1.2.0 with default parameters. Three *Salinicoccus* strains were used as outgroup to root the tree: *Salinicoccus sediminis* SV-16, *Salinicoccus cyprini* CT19, and *Salinicoccus roseus* W12.

### Bacterial strains

A collection of 53 reference strains of validly published staphylococci until 2020 and sensitive to gentamicin has been used, each of them representing one staphylococcal species (S3 Table, sheet Staphylococcal strains). The strain *S. aureus* 8325-4 has been used as a positive control for each experiment, knowing the ability of this strain to adhere to fibronectin-coated surfaces [62]. The strain *S. aureus* DU5883, an isogenic mutant of the *S. aureus* 8325-4 strain not expressing the FnBPs, has been used as a negative control for the fibronectin adhesion assays [62]. The *S. intermedius* strains used for the internalization into osteoblasts and for fibronectin adhesion assays are listed in S3 Table, sheet *S. intermedius* strains.

### Cell culture

Three distinct osteoblast cell lines were cultured: the human osteoblastic MG-63 cell line and two murine osteoblastic cell lines [63,64]. These cells line come from calvaria of transgenic mice: OB-β1^+/+^ expressing a functional integrin β1 subunit and OB-β1^-/-^ deleted for the *itgb1* gene coding for the integrin β1 subunit [64]. Cells were cultivated into 250mL flasks with 11mL of Dulbecco’s modified medium with phenol red, supplemented with 10% of decomplemented fetal bovine serum and 100µg/mL of penicillin and streptomycin. Cells were incubated during one week at 37°C and 5% CO_2_ before plating.

### Bacterial culture

Bacteria were cultivated in 25mL of brain heart infusion broth for 18h at 37°C and 180rpm stirring. Bacterial cultures were centrifugated at 2000*g* for 7min. The pellet was resuspended with 5mL of cell culture medium lacking phenol red. Dilution were plated on tryptone soy agar (TSA) to measure bacterial concentration after overnight incubation at 37°C. Until the infection, the suspension was conserved at 4°C.

### Fibronectin adhesion assays

Fibronectin adhesion assays were conducted in 96 wells flat bottom microplates. The wells were coated with 200µL of 50µg/mL human fibronectin or with 200µL 1% bovine serum albumin (BSA) diluted in phosphate-buffered salin, during 18h at 4°C under constant stirring of 50 rpm. They were washed three times with 200µL of PBS/BSA during 20min at 37°C. The reagent A Baclight^TM^ RedoxSensor^TM^ Green Vitality Life Technologies was distributed in every bacterial suspension of 10^8^CFU/mL to reach a 1µM final concentration. Suspensions were incubated in the dark for 15min at 37°C under constant stirring of 80rpm. A volume of 100µL of each strain was distributed in the fibronectin-coated plate. The microplate was incubated for 45min at 37°C with 80rpm constant stirring. The plate was washed three time with PBS and the last washing PBS was kept for the fluorescent measurement through a plate reader Infinite M Nano+ Tecan. The results are expressed as the percentage of the fluorescence measured for the strain of interest compared to the one of the positive control *S. aureus* 8325-4 after deducing the fluorescence measured for the non-specific interactions with BSA.

### Determination of the internalization capacity of staphylococcal strains

Cells were seeded to obtain 100 000 cells per well after 48h of incubation at 37°C and 5% CO_2_ in 24 wells culture treated plate. The plate was washed two times with 700µL of PBS and cells were infected with 1mL of bacteria diluted in cell culture medium for a multiplicity of infection (MOI) of a hundred bacteria per cell. The number of cells was counted the day of infection, and the real MOI was checked for each experiment. The infected plate was incubated during 2h at 37°C and 5% CO_2_. The medium was removed, and wells were washed one time with PBS. Then, 1mL of 20 µg/mL gentamicin was added in each well. The plate was incubated during 1h at 37°C and 5% CO_2_. After washing with 700µL of PBS, 1 mL of sterile water was added in each well to lyse the cells, the plate was incubated for 30min at 37°C and 5% CO_2_. Lysates were plated of TSA and incubated for 24h at 37°C.

### Determination of the role of B1 integrin in highly internalized staphylococci

The same method as for the determination of the internalization capacity of staphylococcal strains was used. However, the two cell lines OB-β1^+/+^, expressing a functional integrin β1 subunit, and OB-β1^-/-^, deleted for the *itgb1* gene coding for the integrin β1 subunit [64], were plated on the same 24 well culture treated plate for a better comparison of the obtained data.

### Identification of *fnb* genes homologues and analysis of their genomic environment

Blastp v2.13.0+ was used to search *fnb*-like sequences in every genome. The query sequence used for the blastp was the FnBPA reference protein sequence in Uniprot (ID P14738). Only sequences with an identity higher than 25% and a coverage higher than 40% were kept. Interproscan website v92.0 or higher was used to analyze protein domains of all candidates, which were then compared to the reference sequence. Four main domains compose the reference sequence: YSIRK Gram-positive signal peptide (IPR005877), SDR-like Ig domain (IPR041171), Fibrinogen-binding domain 2 (IPR011266) and LPXTG cell wall anchor domain (IPR019931). Protein sequences containing SD repeats were removed as these repeats are characteristic of Sdr proteins [35]. The schematic representation of each protein sequence was constructed using the Interpro domain search software and the SignalP 5.0 signal peptide search software. Then, genomic environments of each candidate were manually investigated to search for other FnBP-like candidates by comparing the environments. An environment is defined as 10 genes: five upstream and five downstream of the FnBP homologous protein (S2 Table).

### Molecular phylogenetic analysis

An alignment of sequences of the 29 homologous proteins was performed using the MAFFT version 7 program with default parameters. Based on this alignment, a tree was constructed using the Neighbor joining method, taking into account all sequences without gaps. The Jones, Taylor and Thornton (JTT) sequence evolution model was applied for tree construction with estimated sequence heterogeneity [65].

## ACKNOWLEDGMENTS

The authors thank the engineers and biologists of the French National Reference Centr for Staphylococci for their advice and expertise. The sequencing of several genomes was performed at the GENomique EPIdémioloique des maladies Infectieuses (GenepII) platform of Hospices Civils de Lyon.

## SUPPORTING INFORMATION CAPTIONS

**S1 Figure. Domain organizations of the 29 identified FnBP-like proteins.** These representations were constructed using the Interpro domain and the SignalP 5.0 signal peptide softwares. Proteins have been ordered according to the species phylogenetic tree and in the order of appearance in genetic environments.

**S2 Figure. Phylogenetic tree of FnBP-like proteins.** This phylogenetic tree was done using MAFFT version 7 program and the Neighbor joining method. A: Phylogenetic tree based on domain A of the FnBP-like proteins. B: Phylogenetic tree based on the entire sequence of FnBP-like proteins.

**S1 Table. Genetic environment of FnBP-like coding genes.** Table summarizing clusters, species, strains, and environmental identifications.

**S2 Table. FnBP-like proteins.** FnBP-like protein candidates and all the conserved FnBP-like proteins. Table detailing FnBP-like proteins, including species, gene identifiers, and associated environments.

**S3 Table. List of all strains studied.** Table of staphylococcal strains, including species, strains, NCBI accessions, and origins.

